# Continuous Daily Head and Trunk Acceleration Recording for the Assessment of Bone Stimulation in Astronauts

**DOI:** 10.1101/2022.01.03.474794

**Authors:** Yoshino Sugita, Gilles Clément

**Author notes:** Corresponding author: Gilles Clément, UNICAEN-INSERM U1075 COMETE University of Caen Normandy, Esplanade de la Paix, 14032 Caen Cedex, France.

## Abstract

The acceleration of the head and hip along the x-, y-, and z-axis of 14 healthy subjects was recorded during two sessions of 12 consecutive hours. The magnitude, frequency content, and root mean square of the acceleration signals were used to determine the type of physical activity (sitting, standing, walking, etc.) during normal daily life on Earth. The acceleration signal slope (jerk) was also calculated to assess whether these activities were sufficient to maintain bone mineral density. These measurements indicated that the changes in vertical acceleration experienced by our subjects during normal daily life were presumably sufficient to maintain bone mineral density. However, these changes might not be sufficient for postmenopausal women and astronauts during long-term exposure to weightlessness during spaceflight.

## INTRODUCTION

On Earth, the measurement of body acceleration has been used as a low cost and power electronic sensor in clinical and home environment to monitor physical activities. Accelerometers have been used to study people with abnormal movement [Burchfield et al. 2007], back-pain [Bussmanna et al. 1998], Parkinson’s disease [Baukje et al. 2010, LeMoyne et al. 2009], sleep disorder [Ojeda et al. 2010], obesity [Cooper et al. 2000], stroke, migraine [Tulen et al. 2000], and elderly person’s fall detection/prevention [Kang et al. 2010, Chen et al. 2010, Culhane et al. 2004, Marschollek et al. 2008].

Heikkinen et al. [2007] showed that vertical body acceleration achieved by drop jumps from a height of 61 cm could increase bone mineral density in postmenopausal women. These authors also proposed that how quickly this acceleration is applied to bones (i.e. the acceleration signal slope, or jerk) is one key factor to maintain bone mineral density. Since astronauts experience loss in bone density during long-duration space flight, we proposed to continuously monitor the acceleration of the head and hip of the astronauts during their daily activity on board the International Space Station (ISS) [Clément et al. 2020]. Computing the acceleration signal slope would help determine if the exercise regimens used by the astronauts in orbit are generating enough acceleration to the antigravity bones for mitigating bone mineral density loss during spaceflight.

The objective of the present study was to measure head and hip accelerations during normal daily life in university students and staffs. Measurements took place during two 12-hour sessions to determine whether test/retest variability. A motion analysis algorithm was used to identify various subject activities (e.g., sitting, walking, climbing stairs, etc.). Based on Heikkinen et al. [2007] model, the acceleration signal slope was also calculated to assess whether these activities were sufficient to maintain bone mineral density. It was expected that the results of this experiment would be helpful to determine the requirements for designing the experiment to be performed on the astronauts on board the ISS.

## MATERIALS AND METHODS

### Participants

Fourteen healthy subjects (7 female, 7 male; aged 23-55) with no history of vestibular or gait abnormalities participated in the experiment. The study was approved by the International Space University Review board and all subjects gave a signed informed consent prior to participating in the study.

### Accelerometers

Two battery-powered, 3-axis inertial measurement units or accelerometers (Gulf Coast Data Concepts, LLC, Waveland, MS, USA) were used to monitor the subject’s linear accelerations. One was attached to a fitted baseball cap, and the other was attached to a waist belt. Miniature laser pointers were fixed on top on both accelerometers. The total weight of the accelerometer, battery, and laser pointer was 55 grams. During calibration sessions, which took place at the beginning of the test session and about 3 to 5 times during the recording, the subject faced a calibration paper sheet and pressed the laser pointer to check that the position of the accelerometers had not changed. For this reason, the accelerometers were placed on the left side of the baseball cap and belt, for the laser beam to point in front of the subject. In previous experiments, other authors had placed one accelerometer at the back of the head [Kavanagh et al. 2008, Oman et al. 1986, MacDougall and Moore 2005], the front of the head [Pelham et al. 2006], or the top of the head [Watt et al. 2003]. Comparison of the results across these studies shows little inconstancies because the centripetal acceleration generated by head movements at the head rim is much smaller that the linear acceleration generated by body physical activity.

Previous studies had also used single accelerometer mounted at waist or sternum for the recognition of some basic activities, such as walking, running, and lying down [Mathie et al. 2004, Sekine et al. 2006, Karantonis et al. 2004, Lee et al. 2003, Khan et al. 2010] In the present experiment, two accelerometers were used to record the accelerations of both the head and hip. Knowing head and hip movements are essential to understand astronauts’ activities under a microgravity environment. Accelerations were measured along the x-axis (forward-backward), y-axis (side to side), and z-axis (up-down) at a rate of 20 Hz. Previous studies indicated that a frequency above 0.45 Hz was critical for physical activity recognition [MacDougall and Moore 2005]. The accuracy of the accelerometers was ±0.004 g, ±0.004 g and ±0.006 g, in the x-, y-, and z-axis respectively.

### Experimental Protocol

The data collection started between 8:00 am and 10:30 am and ended 12 hours later. During the first calibration session, a bubble lever was used to level the accelerometers, and the laser pointer was use for marking the projection of the head and hip accelerometers on a calibration paper sheet. The subjects were asked to come and check the position of the accelerometers regularly during the recording session. During data recording the subjects kept a log file of their activity and wrote down the periods where they were sitting, walking, eating, etc. Each subject was tested during two sessions separated by several days to check the test-retest variability in the responses. After the testing the accelerometers data was transferred to a PC via a USB connector. Data was sorted in a spreadsheet for off-line analysis.

### Data Analysis

First, a Fourier transform was used to measure the frequency content and amplitude of acceleration. Equation (1) was used to find the value of the linear acceleration component along a particular axis. The interval is the number of data points per set, e.g. 40 in this experiment.

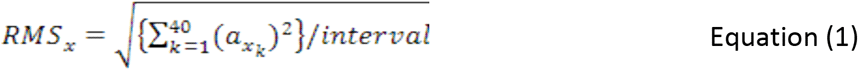

Next, we used the method defined by MacDougall and Moore [2005] based on the frequency content of the z-axis acceleration signal to determine the subject’s locomotion versus non-locomotion. Acceleration data were divided into 60-second interval, and the instantaneous frequency of z-axis acceleration was calculated for each interval. The first step to identify physical activities is to know whether a subject is in locomotion or non-locomotion. Locomotion was defined as the periods where this frequency ranged from 1.5 Hz to 2.5 Hz, whereas non-locomotion is defined as the periods where this frequency was below 1.5 Hz. Once an activity is identified as locomotion, root mean square (RMS) of the acceleration signal was used to identify a subject is walking, running, and falling down (Figure 1).

**Figure 1.**
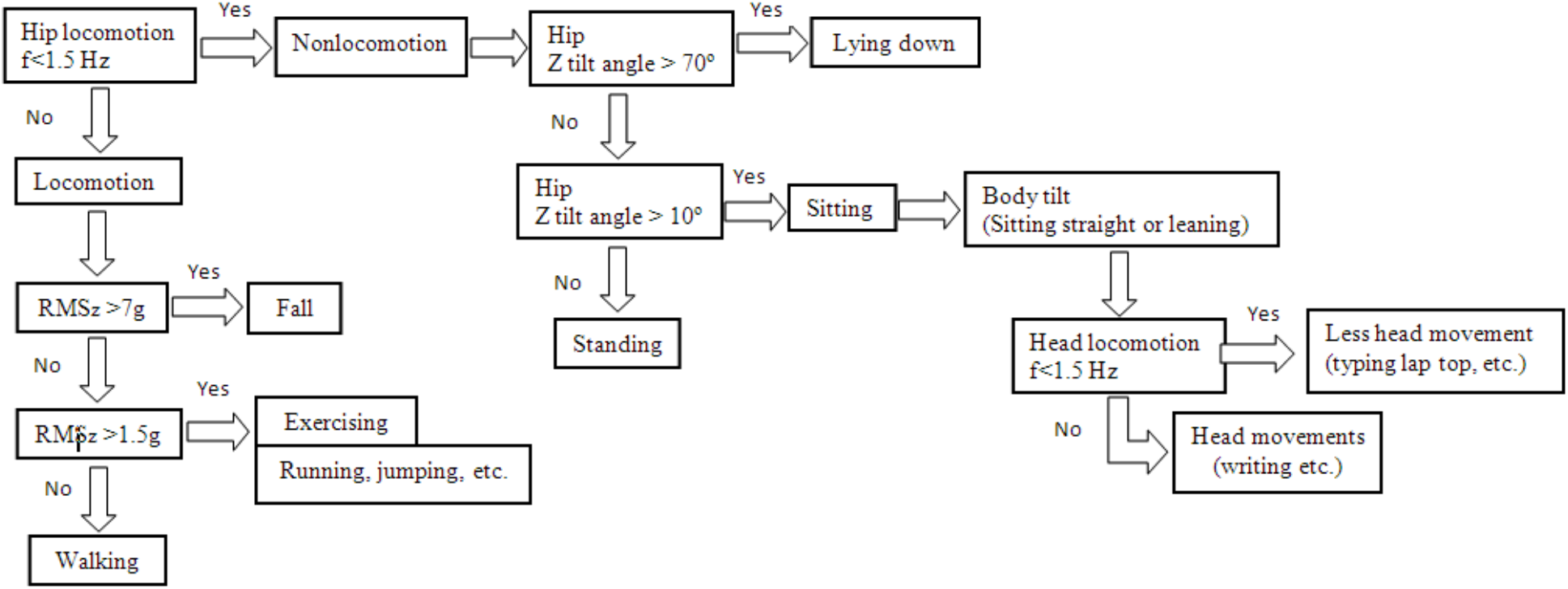
Block diagram used for the activity recognition method.

During the non-locomotion periods, the tilt angles about the yaw, pitch, and roll were calculated by Equation (2), based on the acceleration signals along the x-, y-, and z-axis, respectively.

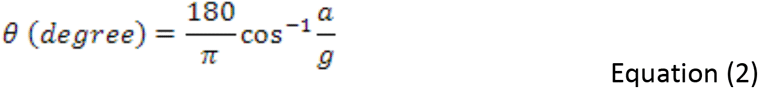

Finally, based on Heikkinen et al. [2007] model, we also calculated the acceleration signal slope when the acceleration was higher than 0.3 g.

## RESULTS

The power spectrum of acceleration along the x-, y-, and z-axis for the 14 individual subjects during the first of their 12-hour recording sessions is shown in Figure 2. Data is very comparable across subjects, despite differences in age, gender, or nationality. Paired t-tests indicated that there was no significant difference between the first and second recording sessions. Larger accelerations are seen along the z-axis, compared with the x- and y-axis, with peaks of frequency at about 2 and 3 Hz. The acceleration along the y-axis (side-to-side) peaked at about 0.8 Hz (which is presumably the centripetal acceleration due to head turn in yaw). Except for one subject, the acceleration along the x-axis was below 0.1 g.

**Figure 2.**
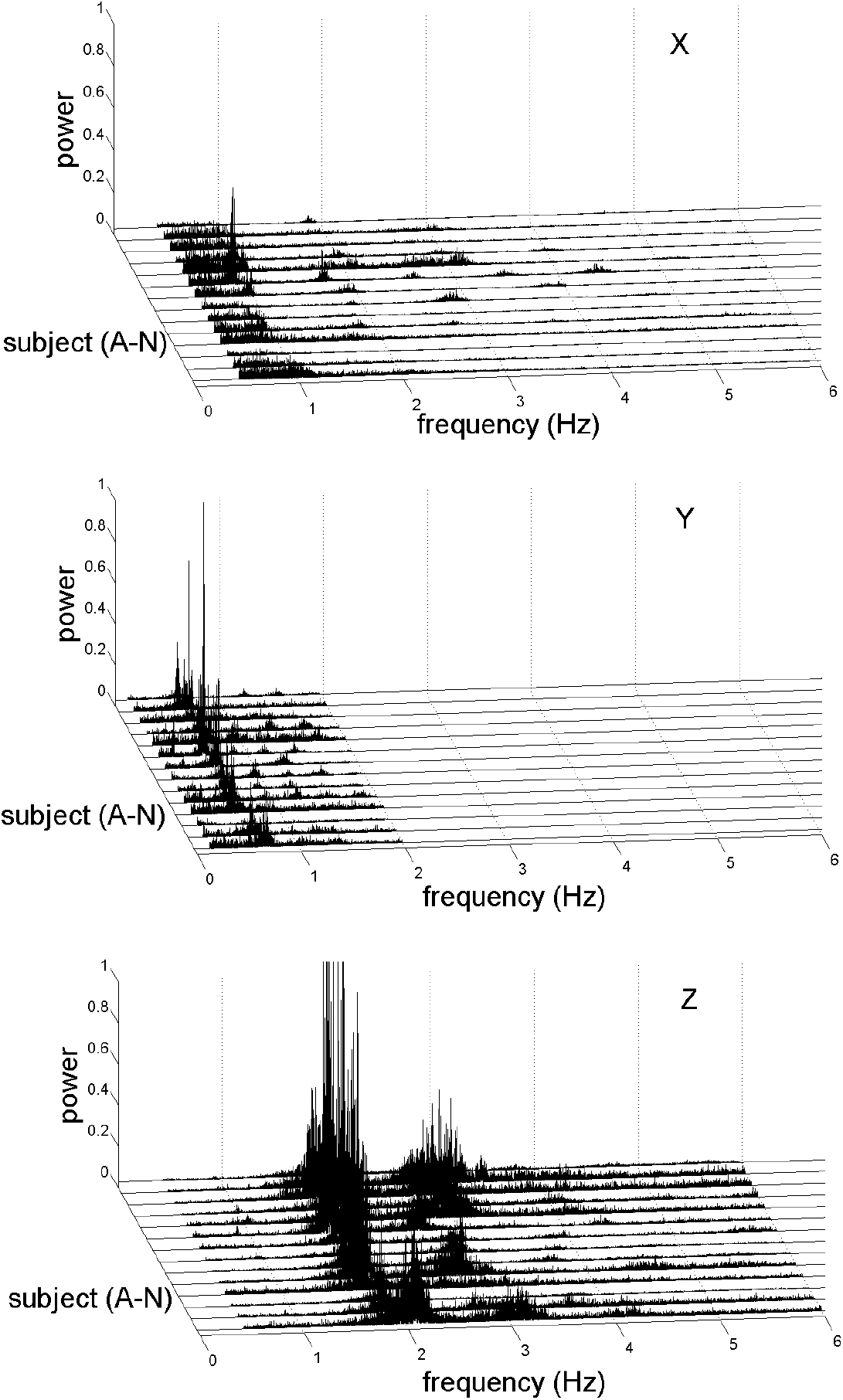
Power spectrum of acceleration along the x-, y-, and z-axis for the 14 individual subjects during a 12-hour recording session.

The acceleration along the z-axis during 12-hour for a typical subject is shown in Figure 3A and B, for the head and hip respectively. The periods where the subject indicated he was walking, sitting, or eating are also shown. Comparison between the signals from the two accelerometers indicates that some activities sometimes generate acceleration at the head only (e.g., during sitting), at the hip only (e.g., between 4.5 and 5 hours), or both. For example, the arrows at 0.5, 3, 8.8, 9.5 hours shows that the vertical acceleration at the hip was below 1 g during these events. It is likely that these accelerations are not vertical body translation, but rather body tilt relative to gravity. The direction of the acceleration suggests that in fact the subject was leaning backward, which is a common posture for students attending a lecture.

**Figure 3.**
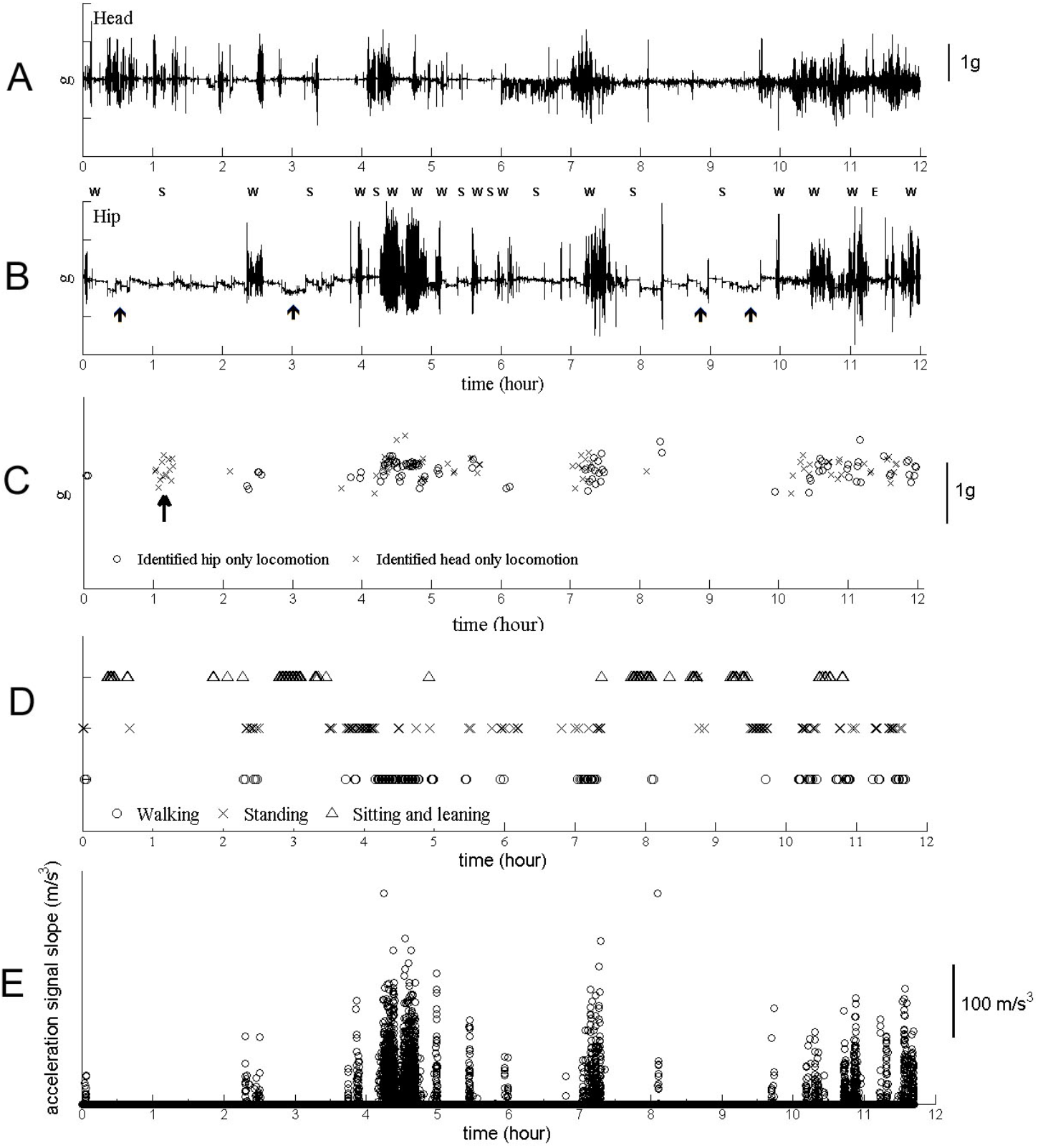
**A,B.** Magnitude of acceleration along the z-axis at the head (A) and the hip (B) for a 23-year old male student. Note: W: Walking, S: Sitting, E: Eating. The arrows point to the times where the subject was presumably tilted relative to gravity. **C.** Symbols indicate when locomotion was identified at the hip only or at the head only using the block diagram defined in Figure 1. **D.** Symbols indicate physical activities (walking, standing, and sitting) also identified by the block diagram in Figure 1. **E.** Z-Axis acceleration signal slope (jerk) for the hip of the same subject, based on the method proposed by Heikkinen et al. [2007]. Each open symbol represents the acceleration slope value at a given time. The jerks that are less than 0.3 g and the negative slope values are not shown.

Based on their frequency content, the acceleration data were used to determine if the subject was in motion (locomotion) or not (non-locomotion). Non-locomotion behavior was defined when frequency was less than 1.5 Hz, and locomotion behavior was when frequency ranged from 1.5 to 2.5 Hz [McDougall and Moore 2005]. As an example, Figure 3C shows when locomotion was identified at the hip only or at the head only, whereas Figure 3D shows three physical activities (walking, standing, and sitting) identified by the block diagram shown in Figure 1. Figure 3E shows the acceleration signal slope for the z-axis at the hip based on the method proposed by Heikkinen et al. [2007]. The maximum value does not exceed 500 m/s^3^.

If a subject’s activity is identified as locomotion, RMS in the z-axis is used to identify falling, walking and exercising. RMS along the z-axis (RMSz) was the smallest among other RMSs (Table 1), which confirms that the major direction of the acceleration was along the z-axis, which is the direction of gravity in the normal life. If the subjects experience higher acceleration activities, such as drop jumping, then the percentage of RMSz should increase. The higher RMSy value compared to RMSx is primarily due to the direction of walking (i.e. walking) and leaning forward/backward when sitting.

**Table 1.** Comparison of acceleration changes (RMS) along the x-, y-, and z-axis for the head and hip during the first 12-hour recording session in all 14 subjects. Mean (SD).

|  | Head (%) | Hip (%) |
| --- | --- | --- |
| RMSx | 14.3 (SD 14.7) | 13.8 (SD 14.7) |
| RMSy | 37.8 (SD 17.6) | 46.8 (SD 17.6) |
| RMSz | 9.4e-3 (SD 2.8e-2), | 2.3e-2 (SD 2.8e-2) |

As shown in Table 2, most of the 14 subjects’ activity fell onto “non-locomotion” behavior. Subject 13 had the highest percentage for the head and hip movements (21.6 and 21.8%, respectively). This subject is a teacher’s assistant who often noted in his log that he walked around in the building to prepare equipment and classrooms. The other subjects were students and faculty. In average, our subjects expressed the same amount of head locomotion percentage than showed in previous studies [MacDougall et al. 2005]. At the end of the recording session, subjects were asked to guess the duration for which they had any locomotion behavior, or exercise, during their recording session. It is interesting to note that all subjects overestimated their amount of daily exercise (33% for subjective evaluation compared to 19% for acceleration measurements, in average).

**Table 2.** Percentage of the overall recording period where the frequency of the z-axis acceleration ranged from 1.5-2.5 Hz, suggesting that the subjects had a locomotion behavior. Measured performed during the first 12-hour recording session in all 14 subjects, using both the head and hip acceleration data. The column on the right shows the percentage of the overall recording session that the subjects verbally indicated as locomotion.

| Subject | Head (%) | Hip (%) | Locomotion (%) |
| --- | --- | --- | --- |
| 1 | 16.5 | 19.3 | 25 |
| 2 | 10.5 | 10.9 | 20 |
| 3 | 2.2 | 2.0 | 10 |
| 4 | 18.8 | 8.1 | 60 |
| 5 | 2.9 | 20.1 | 35 |
| 6 | 12.3 | 11.8 | 30 |
| 7 | 8.5 | 8.6 | 20 |
| 8 | 10.2 | 11.6 | 20 |
| 9 | 13.8 | 14.9 | 60 |
| 10 | 11.1 | 12.9 | 40 |
| 11 | 10.7 | 13.3 | 40 |
| 12 | 15.1 | 21.2 | 65 |
| 13 | 21.6 | 21.8 | 20 |
| 14 | 10.2 | 10.2 | 20 |

Tilt of the body in yaw, pitch, and roll based on the acceleration signals during non-locomotion behavior is shown in Figure 4A, B, C. The tilt angle in the y- and z-axis shows the posture on the y-z plane. Zero degree in the y- and z-axis means the subject is standing straight. The high tile angle in the y- and z-axis shows the subject is lying down. The tilt angle in the x-axis shows the posture on the x-z plane. Zero degree in the x-axis means the subject is lying down and high degree near 90 degrees mean that the subject is standing or sitting straight. The information above is useful to identify physical activities. The tilt angle in Figure 4D shows a high correlation (R^2^ = 0.99) between 2 tilt angles using the trigonometry relationship between gravity direction, y- and z-axis.

**Figure 4.**
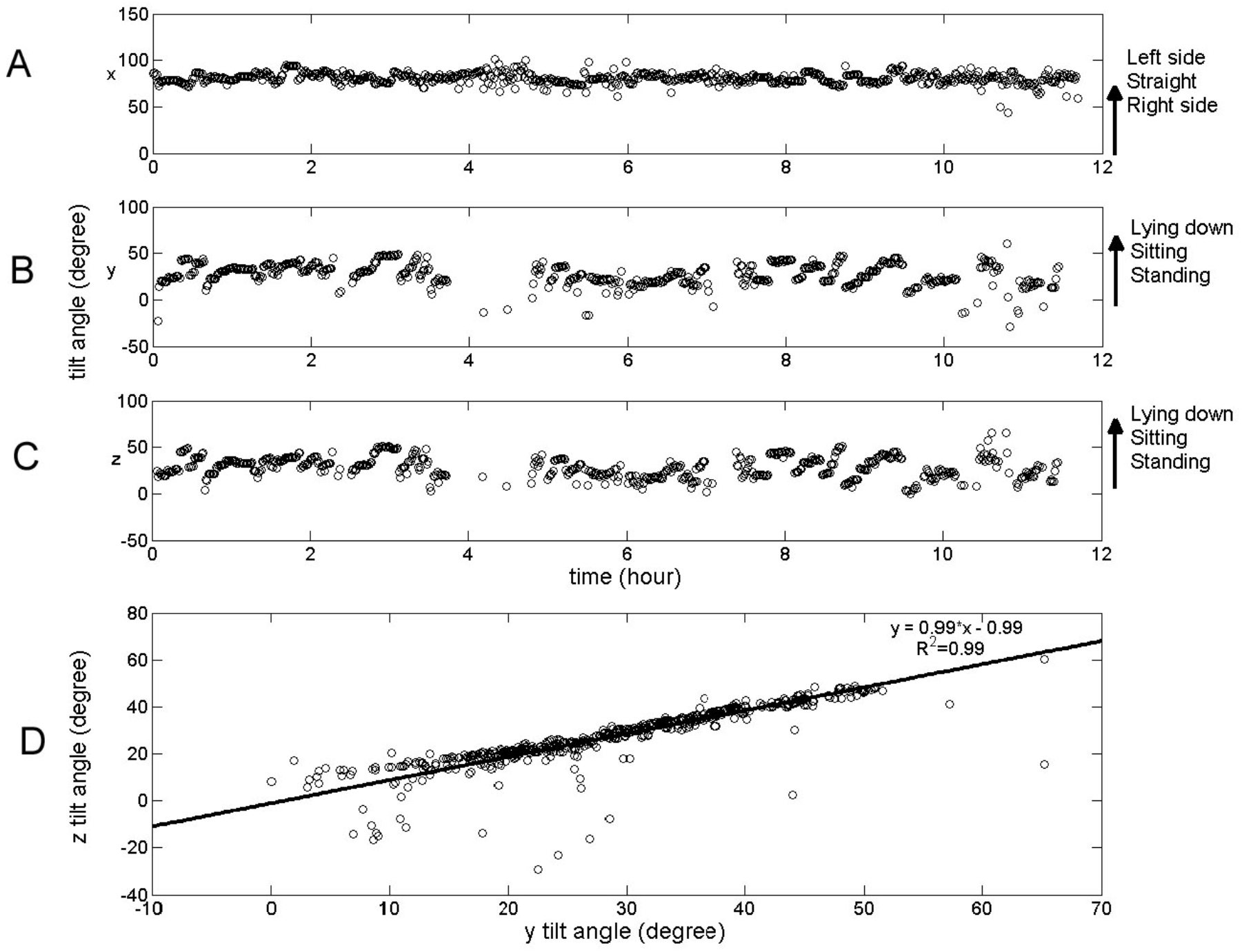
**A,B,C.** Tilt angle of the body in yaw (A), pitch (B), and roll (C) based on the hip acceleration signals along the x-, y-, and z-axis, respectively. **D.** Correlation between the body tilt angles relative to gravity calculated from the acceleration along the x- and y-axis during nonlocomotion.

## DISCUSSION

Our results show that a motion analysis based on the magnitude, frequency content, and RMS of linear accelerations along the x-, y-, z-axis of the head and hip can be successfully used to determine various types of physical activity (e.g. sitting, standing, walking), including locomotion and non-locomotion patterns during normal daily life.

It has been proposed that the change in acceleration (acceleration signal slope) could be used to determine whether this physical activity is sufficient to mitigate bone loss. Judex et al. [2000a] showed that drop jump produced large peak strain rates (+740%) compared with baseline walking. They also showed that treadmill running strain peak was 36% higher than walking strain peak, but did not enhance bone growth [Judex et al. 2000b]. Heikkinen et al. [2007] measured the acceleration slope values for different exercise patterns, such as drop jump, running, walking and stepping in women aged 20-58 years. They also compared the acceleration slope values between two groups of subjects: a control group (n=30) and an exercise group (n=34), and they measured bone mineral density in all the subjects before the experiment and after 12 months. They determined that the osteogenic threshold was 1000 m/s^3^, which could be achieved by drop jumping and running but not in walking or stepping. The acceleration signal slope in our subjects during stepping and walking was less than 500 m/s^3^. This exercise level may be sufficient for a 23-35 healthy subjects, but may not be sufficient for the postmenopausal women discussed in the Heikkinen’s paper, or in astronauts during long-duration spaceflight.

The deconditioning of astronauts’ health during long space missions is a well-recognized problem. The deconditioning includes bone density, reduced muscle strength, exercise capacity, motion sickness and orthostatic tolerance. Countermeasures for mitigating this deconditioning play an important role in space mission success. Current in-flight countermeasures for bone and muscle strength loss include exercise utilizing load-bearing treadmills, resistive exercise devices, and cycle ergometers [Clément & Reschke 2008]. For treadmill exercise, elastic straps between the subject’s shoulder and the treadmill generate the mechanical g-load. However, the straps are uncomfortable and as a result, the subjects do no tight them enough. Monitoring the acceleration at the hip would help assess the duration and magnitude of mechanical load exerted on the longitudinal body axis during exercise on board the ISS.

Sensors have been used in space to monitor astronauts’ core body or skin temperature, heart rate, and radiation exposure, as well as EKG, O2 consumption and CO2 consumption during extravehicular activity (EVA) activity [Clément 2011]. Smart watches continuously worn by astronauts have also been used to provide sleep schedule variability, sleep quantity, and sleep quality statistics. However, these devices provide actigraphy data and do not differentiate between acceleration across the x-, y-, and z-axis.

Continuous head and hip acceleration monitoring could also be helpful for mitigating orthostatic intolerance and motion sickness. Bed rest studies have shown that aerobic exercise training helped maintain plasma volume and work capacity [Greenleaf et al. 1994]. However, aerobic exercise alone was not effective in increasing tolerance to post-bed rest orthostatic challenges. Some studies have shown that short, intermittent exposure to 1 g along the z-axis significantly improved orthostatic tolerance after prolonged head-down bed rest [Vernikos et al. 1996; Xiao et al. 2005, Lathers et al. 1991, Muir et al. 2011]. The orthostatic impairment observed in astronauts on return from space travel closely resembles the clinical syndrome of orthostatic intolerance after bed rest [Saltin 1992]. In addition, space motion sickness during transitions between gravitational states, such as during entry into weightlessness and immediately after landing, is generated by head movements in pitch and roll [Oman et al. 1986]. Monitoring physical activities of astronauts by accelerometers has been proposed to enhance exercise methods that would prevent orthostatic symptoms and motion sickness [Cox et al. 2002, Ertl et al. 2002, Bloomberg et al. 2015].

During this ground-based study, the major complains from our subjects was related to the weight of the head accelerometer and the baseball cap which was uncomfortable for 12-hour use. A better design of the head strap and lightweight accelerometers are required for the study on the astronauts. Wearable accelerometers can provide valuable information on characterizing individual daily physical activities, including locomotion or non-locomotion behavior, and head tilt relative to gravity. This information could be used for defining better individual prescription or countermeasures against the effects of inactivity on long-duration spaceflight, bed rest, or sedentary life.

